# How tree diversity and ectomycorrhizal dominance affect biomass allocation of mixed deciduous forests

**DOI:** 10.64898/2026.01.29.702198

**Authors:** Annalena Ritter, Daniela Yaffar, Ina C. Meier

## Abstract

Biomass and surface area allocation affect resource uptake and carbon (C) residence time in forests, but the influence of tree diversity on allocation remains poorly understood. Moreover, mycorrhizal associations can alter this relationship, which has been rarely tested in mature forests. We investigated the role of both the proportion of ectomycorrhizal (ECM) trees and tree diversity on tree biomass and surface area allocation across a dual gradient of tree diversity (0 – 1.68 Shannon diversity) and ECM dominance (0 – 100 %) in a mixed deciduous forest area in Central Germany. We found that the two gradients affected tree biomass and surface area differently and mostly independently. Tree diversity had no significant effect on biomass or surface area in the investigated forest area, but increased the spatial variability of the leaf area index (LAI) from 21 % to 40 %. In contrast, a higher proportion of ECM trees was associated with an increase in fruit biomass (from 10 to 141 g m^-2^) and LAI (from 4 to 7 m^2^ m^-2^). Although tree diversity and the portion of ECM produced similar parsimonious models for explaining belowground biomass and surface area, neither showed a significant direct effect. Notably, their interaction enhanced the spatial variability of fine root biomass and root surface area; that is, forests with high diversity and a greater proportion of ECM trees exhibited a more heterogeneous distribution of fine roots. Allocation to fine root biomass appeared independent of tree diversity and the proportion of ECM trees, being influenced primarily by stand structure, with higher allocations observed in stands with lower stem basal area. We conclude that biomass allocation in this Central European Forest, where resource availability is relatively uniform, is primarily productivity-driven. A comparison of the biotic influences shows that ECM trees have a stronger control on aboveground surface area and fruit biomass than tree diversity, which may contribute to the ability of dominant ECM trees, such as European beech, to outcompete light competitors, but also puts temperate ECM forests at risk of physiological failures in increasingly drier future conditions.

## Introduction

Global forests account for 80 % of Earth’s total plant biomass and contain more carbon (C) in their ecosystem than is stored in the atmosphere (Pan et al. 2013). This enormous biomass can be allocated aboveground or belowground. Such biomass allocation to different organs influences not only tree C and nutrient uptake capacities and optimizes tree growth, but it also determines soil C residence time and terrestrial C storage as a consequence (Mokany et al. 2006), since root litter decomposes more slowly than leaf litter (Freschet et al. 2013; Gholz et al. 2000) leading to more stable soil C forms such as mineral-associated organic matter (MAOM; Prescott 2010). It has already been well established that plants allocate biomass optimally in response to the abiotic environment to enhance uptake of the most limiting resource (optimal partitioning theory; Bloom et al. 1985). This fundamental theoretical framework applies to plants in many environments and can be used to predict allocational shifts in response to climate change. However, this framework has recently been challenged by biotic influences altering the relationship between allocation and environmental response (e.g., Rong Liu et al. 2021; Puglielli et al. 2021). Particularly, it has been shown that the symbiotic interaction with mycorrhizal fungi influences biomass allocation strategies in a warming world (Zhou et al. 2022). Improving our understanding of biotic influences on biomass allocation will be essential to understanding the consequences of global species loss for soil C stabilization.

The majority of trees form symbioses with mycorrhizal fungi (Wang and Qiu 2006). Mycorrhizal symbiosis can affect biomass allocation of trees due to altered resource uptake capacity and nutrient foraging strategies of mycorrhizal root systems (Zhou et al. 2022). Mycorrhizal hyphae can replace, to some extent, the function of fine roots by increasing their surface area and mobilizing recalcitrant nutrient sources in exchange for C supplied by the root host (van der Heijden et al. 2015; Bergmann et al. 2020), which can decrease the C investment into root biomass as a consequence. The most common mycorrhizal types in trees are arbuscular mycorrhizal (AM) and ectomycorrhizal (ECM) associations. Since most tree genera are predominantly associated with only one of the two major mycorrhizal types (Brundrett and Tedersoo 2020), the mycorrhizal association can be used as a functional grouping of tree species into groups of similar environmental response. The two mycorrhizal types differ fundamentally in their nutrient foraging strategy and nutrient economy, where AM trees dominate in nutrient-rich environments with faster biogeochemical cycling, and ECM trees in nutrient-poorer environments in which biogeochemical cycles are slowed down (Phillips et al. 2013). According to optimal partitioning theory, this should lead to smaller root:shoot ratios in AM trees than in ECM trees as a consequence of less biomass being allocated to roots in ecosystems where decay rates are fast and belowground resources are less limiting. Estimations of the global root:shoot ratios of AM and ECM trees did not show significant differences between the two mycorrhizal associations, even though a tendency towards smaller root:shoot ratios in AM trees was visible (Barceló et al. 2023). By contrast, AM trees had higher root:shoot C pools than ECM trees in temperate forests of Northeast China (Wang et al. 2023). Jevon and Lang (2022) suggested that AM trees allocate globally 4 % more biomass to roots, and that this effect is greater than that of climate. Maintenance costs differ between AM and ECM root systems, and lower C investment in the AM association can lead to greater root biomass and belowground overrepresentation of AM tree species compared to ECM tree species (Valverde-Barrantes et al. 2018). Particularly in the AM association, roots provide intraradical habitat for their fungal partners by increasing the root cortical area (Bergmann et al. 2020), which may add to greater root biomass in AM trees. However, it remains unclear whether root maintenance costs (host-symbiont interaction) and/or the prevailing nutrient economy (resource availability) influence the biomass allocation of mycorrhizal trees.

Biotic interactions with neighboring trees are an additional factor that can alter the within-tree biomass allocation. In tree diversity experiments with young temperate trees or mature tropical trees, tree diversity caused aboveground overyielding and changed the canopy architecture, but did not affect biomass allocation to roots (Guillemot et al. 2020; Martin-Guay et al. 2020). This increase in aboveground biomass can be explained by tree size – that is, it is productivity-driven – since larger, overyielding trees produce relatively more stem biomass to support their structure. Additionally, plasticity-driven changes in crown structure support efficient space-use to decrease competition for light among neighboring trees (Guillemot et al. 2020). While several studies provide support for aboveground (stem) overyielding, discrepancies are reported for the diversity effect on fine root biomass of forest ecosystems (Xu et al. 2020). Fine root biomass was shown to increase with tree diversity at lower stand density, which was a result of root biomass aggregation in the nutrient-rich topsoil and not of greater vertical root partitioning (L.-T. Zheng et al. 2021). Other studies in mature mixed forest stands found no evidence of belowground overyielding (Meinen et al. 2009; Jacob et al. 2013), potentially due to fully occupied topsoil space and the absence of vertical soil segregation into deeper, nutrient-poorer soil layers (i.e., absent niche partitioning). Alternatively, rather than increasing their biomass, roots can become more efficient when soil resources are not limiting, which could lead to no change or even reduced allocation to root biomass in diverse forests, as shown for temperate trees in a common garden tree diversity experiment (Archambault et al. 2019) and in mixed forest stands (Wambsganss et al. 2021). Consequently, tree diversity may act as an amplifier of resource-driven optimal allocation patterns of biomass in forest trees.

In our dual-gradients study, we aimed to analyze the influence of tree diversity and mycorrhizal association type on tree biomass and surface area allocation to leaves, stems, and fine roots in a mature, mixed deciduous forest area with similar resource availability. We hypothesized that (i) tree diversity increases leaf and stem production. We further hypothesized that (ii) ECM-associated trees allocate relatively more biomass and surface area to leaves, and AM-associated trees allocate more biomass and surface area to fine roots due to different environmental responses. Consequently, (iii) the mycorrhizal association type alters the relationship between tree diversity and biomass allocation, where diverse ECM stands have an amplified leaf biomass allocation and diverse AM stands have a diminished allocation response to tree diversity.

## Materials and Methods

### Study sites

We conducted our research in an old-growth mixed forest of the Hainich National Park in Central Germany (51°08’N, 10°51’E), which is, with a size of *c.* 130 km^2^, one of the largest deciduous forest stands in Central Europe (Nationalpark-Verwaltung 2022a). Since 1997, the southern part of the forest has been a national park (Nationalpark-Verwaltung 2022b, 2022a). The forest is located on eutrophic Luvisols (IUSS Working Group WRB 2022), which have developed from a base-rich Pleistocene loess layer over Triassic limestone (European lithostratigraphic unit: Middle Muschelkalk). The mineral soil texture (0 – 30 cm soil depth) is characterized by low sand (< 5 %) and high silt contents (Guckland et al. 2009). The climate is semi-humid with a mean annual temperature of 7.7 °C and a mean annual precipitation of 590 mm.

The studied forest has been unmanaged over the last 40 years and has forest continuity for at least the last 200 years (Schmidt et al. 2009), and therefore represents ancient woodland (Wulf 2003). The vegetation can be classified as Stellario-Carpinetum (starwort-oak-hornbeam forest, interfused with elm trees), with up to 16 tree species co-occurring in this mixed hardwood stand. In our study, these 16 tree species, which are frequently dominant or subdominant trees in the natural forest vegetation of Central Europe and represent two mycorrhizal types (Brundrett and Tedersoo 2020), were included. Arbuscular mycorrhizal (AM) deciduous tree species were represented by common ash (*Fraxinus excelsior* L.), Norway maple (*Acer platanoides* L.), and sycamore maple (*Acer pseudoplatanus* L.), with minor admixtures of wild cherry (*Prunus avium* L.), checker tree (*Torminalis glaberrina* (Gand.) Sennikov & Kurtto), horse chestnut *(Aesculus hippocastanum* L.), and thornapple (*Crataegus spp.*). Ectomycorrhizal (ECM) tree species were mainly represented by European beech (*Fagus sylvatica* L.), pedunculate oak (*Quercus robur* L.), and sessile oak (*Quercus petraea* (Matt.) Liebl.), with minor admixtures of large-leaved lime (*Tilia platyphyllos* Scop.), European hornbeam (*Carpinus betulus* L.), common hazel (*Corylus avellana* L.), and evergreen Norway spruce (*Picea abies* (L.) H. Karst.), and Scotch pine (*Pinus sylvestris* L.). There was also a minor admixture of willow (*Salix spp.*), which can be associated with both mycorrhizal types. The most abundant tree genera according to basal area measurements were AM *Fraxinus* and *Acer*, and ECM *Fagus* and *Quercus*. In this forest, we selected twelve study sites (size: 30 m x 30 m each) with comparable stand structure that represented two biotic gradients (Appendix S1: Figure S1): (1) a tree species diversity gradient ranging from Shannon’s diversity index (H’) of 0.00 to 1.68 (hereafter referred to as Shannon tree diversity) and (2) a mycorrhizal type gradient ranging from 100 % AM trees to 100 % ECM trees (hereafter referred to as proportion of ECM trees; Table 1). Both gradients are based on tree basal area to account for the greater relative influence of larger trees on the forest community. Site selection criteria were (1) closed canopy without major gaps, (2) even-aged stand structure with mature trees, (3) no or minor influence of coniferous trees, (4) similar stem densities, and (5) level to only slightly inclined terrain. The longitude, latitude, and altitude of the center of every site were measured with a global positioning system (GPS; Oregon 700, Garmin, Olathe, USA).

**Table 1:** Topographic, stand structural, and edaphic properties of 12 mature deciduous forest sites along a Shannon tree diversity and ectomycorrhizal (ECM) tree gradient.

| Site # | 1 | 2 | 3 | 4 | 5 | 6 | 7 | 8 | 9 | 10 | 11 | 12 |
| --- | --- | --- | --- | --- | --- | --- | --- | --- | --- | --- | --- | --- |
| East Coordinates | 10°40' | 10°46' | 10°53' | 10°46' | 10°50' | 10°47' | 10°47' | 10°49' | 10°47' | 10°52' | 10°51' | 10°49' |
| Nord Coordinates | 51°10' | 51°10' | 51°08' | 51°07' | 51°08' | 51°11' | 51°07' | 51°08' | 51°08' | 51°09' | 51°09' | 51°08' |
| Altitude (m a.s.l.) | 440 | 360 | 330 | 510 | 400 | 300 | 480 | 390 | 440 | 340 | 390 | 400 |
| Tree species richness (S) | 1 | 1 | 2 | 5 | 5 | 5 | 6 | 7 | 6 | 6 | 8 | 8 |
| Shannon index (H') | 0.00 | 0.00 | 0.56 | 0.72 | 1.11 | 1.22 | 1.31 | 1.37 | 1.44 | 1.48 | 1.49 | 1.68 |
| Evenness (E) | - | - | 0.99 | 0.71 | 0.58 | 0.55 | 0.85 | 0.65 | 0.83 | 0.86 | 0.81 | 0.82 |
| ECM trees [%] | 100 | 100 | 0 | 1 | 71 | 95 | 47 | 94 | 46 | 57 | 11 | 79 |
| *Fagus sylvatica* [%] | 100 | 100 | 0 | 1 | 60 | 30 | 23 | 38 | 30 | 20 | 0 | 40 |
| Basal area [m<sup>2</sup> ha<sup>-1</sup>] | 36 | 31 | 43 | 45 | 34 | 50 | 50 | 43 | 44 | 45 | 26 | 30 |
| Stem density [n ha<sup>-1</sup>] | 333 | 320 | 476 | 613 | 355 | 670 | 357 | 355 | 401 | 401 | 290 | 309 |
| Tree height [m] | 24 | 23 | 22 | 24 | 22 | 22 | 21 | 23 | 25 | 25 | 17 | 22 |
| Tree DBH [cm] | 31 | 28 | 31 | 28 | 31 | 25 | 35 | 36 | 33 | 33 | 30 | 31 |
| Soil pH (CaCl<sub>2</sub>) | 4.4 | 4.7 | 6.6 | 4.8 | 4.7 | 4.2 | 4.8 | 4.3 | 6.1 | 4.7 | 6.4 | 4.4 |
| Soil C/N ratio [g g<sup>-1</sup>] | 11.1 | 11.4 | 10.8 | 12.1 | 11.5 | 12.2 | 11.6 | 11.6 | 12.8 | 11.7 | 11.4 | 12.6 |

The Shannon tree diversity index (H') and the ECM tree proportion (%) are based on the total basal area per tree species.
DBH = diameter at breast height

### Stand structure

In February 2022, all trees with a diameter ≥ 7 cm in the 30 x 30 m sites were counted to determine the number of tree species (n), their evenness, and the stem density (n ha^-1^). The mean diameter at breast height (DBH, 1.3 m height) and stem basal area (m^2^ ha^-1^) were also determined. The summed basal area per tree species was used to estimate the Shannon tree diversity and the proportion of ECM trees. The trees were measured for treetop height with an optical tree height meter (Vertex IV; Vertex, Haglöf, Sweden). Annual diameter growth was measured with dendrometer bands (D1, UMS, Germany) from 2022 to 2024 for ten dominant trees, and the mean annual basal area increment (cm^2^ a^-1^) per mycorrhizal type was calculated (n = 0 to 10). We use basal area increment as a proxy for wood growth and refrain from using allometric functions, as these usually originate from younger trees in monocultures, which have a smaller stem fraction than mature trees in a mixed forest stand (Poorter et al. 2012).

### Aboveground and belowground tree biomass and surface area

Stand leaf and fruit mass (in g m^-2^; equals the annual leaf and fruit production) was determined by litter trapping in 2022. Ten litter buckets (65 cm in diameter) were positioned systematically along two parallel lines at each study site. The traps had holes at the bottom to allow water drainage. All 120 traps were emptied once immediately after the autumnal litter fall. The litter samples were stored at 4 °C for no longer than three weeks and sorted into leaf and fruit fractions of the different tree species. Other litter components, such as twigs, moss, and flowers, were removed. Fifty intact leaves per trap were randomly selected and analyzed for leaf area with WinFOLIA (version 2020a; Régent Instruments, Quebec, Canada). All leaves and fruits were dried (70 °C, 72 h) to obtain dry weight. The leaf area index (LAI; in m^2^ m^-2^) was calculated by multiplying the total leaf mass per ground area by the average specific leaf area (SLA; in m^2^ kg^-1^).

In November and December 2021, fine root biomass (in g m^-2^) was determined in ten soil samples per site, which were taken with a soil corer (3 cm in diameter) from the uppermost 20 cm of mineral soil at random locations. The samples were immediately transported to the laboratory and stored at 4 °C for no longer than four weeks. Roots were washed out of the soil using a sieve with a mesh size of 0.5 mm. Only fine roots with a diameter of under 2 mm were included in the analysis. Fine root segments of trees were picked out by hand and sorted into live and dead roots based on color, structure of the root surface, root elasticity and turgescence, and branching structure. Fine roots were scanned with WinRHIZO (version 2021a; Régent Instruments, Quebec, Canada) to determine their surface area. All fine roots were dried (70 °C, 72 h) and weighed to obtain the fine root biomass of the site (in g m^-2^). The root area index (RAI; in m^2^ m^-2^) was calculated by multiplying root biomass per ground area with the average specific root area (SRA; in m^2^ kg^-1^). We compared the organs for resource uptake by calculating the root fraction (%) from their portion in leaf and fine root biomass, and the root area fraction (%) from their portion in RAI and LAI.

### Soil sampling and chemical analyses

In spring and summer 2022 and 2023, soil samples were taken at four random locations per study site with a soil corer (5 cm in diameter) at 0 – 10 cm of the mineral soil (n = 4 samplings x 4 locations). Roots, organic debris, and stones were removed from soil samples by hand. After air-drying, the soil samples were ground, and the total C and N contents were analyzed by elemental analysis (Vario EL CUBE, Elementar, CHNS mode). The pH of each study site was measured in 0.01 M CaCl_2_ (Seven2Go pH-Messgerät; Mettler Toledo) for three additional soil samples (n = 4 samplings x 3 locations).

### Statistical analyses

Statistical analyses were performed with R Statistical Software version 4.4.2 (R Core Team 2023). Study site means and coefficients of variation (CV) were calculated and used for subsequent analyses. The probability of fit to a normal distribution of the residuals was tested with a qq-plot and homoscedasticity with a residual plot. Significance was determined at p ≤ 0.05, and marginal significance at p ≤ 0.1. The number of compositional, stand structural, and edaphic parameters to be considered was reduced by a principal component analysis (PCA; Table 2). Those principal components explaining ≥ 75 % of the variance were used as independent explanatory variables in regression analyses to study their influence on biomass production, allocation, and surface area aboveground and belowground. In addition, Akaike’s Information Criterion (Sakamoto et al. 1986) corrected for finite sample size (AICc) was used to test which linear model (simple or multiple), with Shannon diversity and/or the proportion of ECM trees as independent variables, best explained biomass and surface area allocation. For this, the “aictab” function from the “AICcmodavg’’ package was used (Mazerolle 2023). Statistical coefficients, statistical significances, and model parameters are depicted in the supplementary materials (Appendix S1: Tables S1 – S5).

**Table 2:** Principal components analysis (PCA) on major compositional, stand structural, and edaphic gradients across the investigated 12 mature deciduous forest sites.

|  | Axis 1 | Axis 2 | Axis 3 |
| --- | --- | --- | --- |
|  | ECM Gradient<sup>1</sup> | Diversity Gradient | Stand Structure |
| Eigenvalue | 1.459 (35.5 %) | 1.292 (27.8 %) | 1.081 (19.5 %) |
| Shannon index (H') | -0.007 | **-0.645** | 0.393 |
| ECM trees [%] | **0.513** | 0.331 | 0.378 |
| Basal area [m<sup>2</sup> ha<sup>-1</sup>] | 0.278 | -0.327 | **-0.603** |
| Tree height [m] | **0.492** | -0.04 | -0.474 |
| Soil pH (CaCl<sub>2</sub>) | **-0.578** | -0.178 | -0.25 |
| Soil C/N ratio [g g<sup>-1</sup>] | 0.288 | **-0.578** | 0.228 |

The most characteristic variables (according to their loading) of each PCA axis are in bold.
ECM = ectomycorrhizal

## Results

The investigated forest sites were located across three environmental gradients: tree diversity, proportion of ECM trees, and stand structure (Table 2). Based on PCA analysis, the main environmental gradient among our study sites, which explained 36 % of the total compositional, structural, and edaphic variability, was composed of soil pH, the proportion of ECM trees, and tree height, where lower soil pH was associated with a higher proportion of ECM trees and higher tree height (PCA axis 1: ECM gradient). The second environmental gradient was a gradient in which Shannon tree diversity and soil C/N ratio explained 28 % of the total variability (negative loadings; PCA axis 2: Diversity gradient). The third environmental gradient was a gradient in which basal area explained 20 % of the total variability (negative loading; PCA axis 3: Stand structure gradient).

According to AICc scores, biomass and surface area were best explained by simple linear models that either included only the proportion of ECM trees or the Shannon tree diversity index, and not by the interaction of both influences. For fruit production and LAI, the best explanatory variable was the proportion of ECM trees. Other biomass and morphological parameters were best explained by both Shannon tree diversity and the proportion of ECM trees, but slightly better by Shannon tree diversity in most cases (with the exception of RAI fraction). The spatial variability of aboveground and belowground biomass and surface area was best explained by simple (CV of stem increment, fruit production, leaf production, and LAI) and multiple linear models (CV of fine root mass and RAI; Appendix S1: Table S2). For the spatial variability of LAI, the best explanatory variable was the Shannon tree diversity index. The spatial variabilities of stem increment, fruit production, and leaf production had similar low AICc for both Shannon tree diversity and the proportion of ECM trees. Still, the CV of fruit and leaf production was slightly better explained by Shannon tree diversity, and the CV of stem increment was slightly better explained by the proportion of ECM trees.

### Tree diversity effect on tree biomass and surface areas

An increase in the Shannon tree diversity index from 0.0 to 1.7 along the investigated gradient had no significant influence on biomass production aboveground and belowground (Appendix S1: Figure S3a). Similarly, tree diversity did not influence the leaf and root surface area indexes (Appendix S1: Figure S2a, b).

The main influence of tree diversity was on the spatial variability of LAI, which significantly increased with tree diversity from 21 % at monospecific sites to 40 % at diverse forest sites (p = 0.006; Table 3).

**Table 3:**
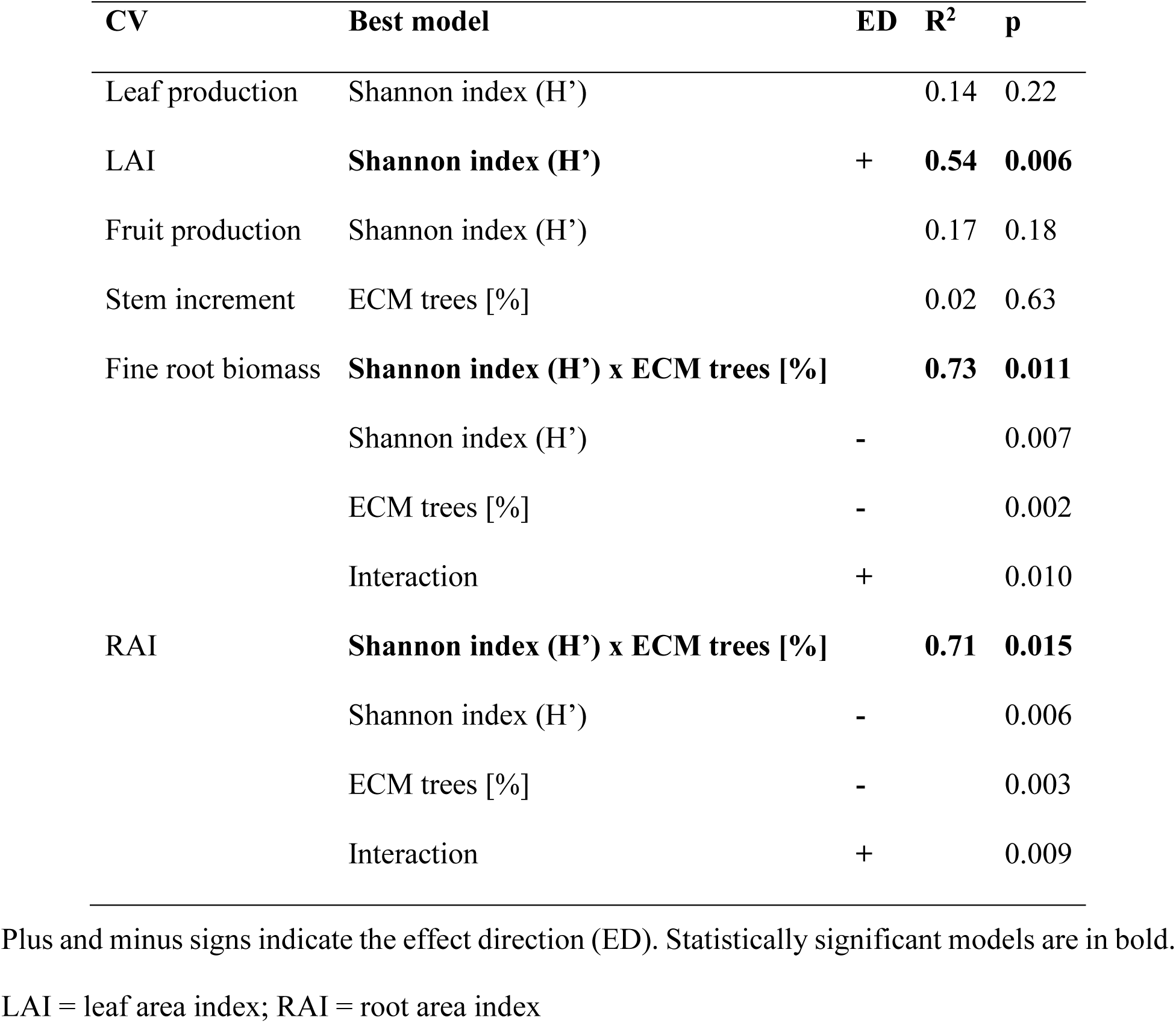
Linear regression analyses on the influence of Shannon tree diversity and ectomycorrhizal (ECM) trees on the coefficients of variation (CV) of aboveground and belowground biomass, production, and morphology of 12 mature deciduous forest sites, utilizing the best model (by AICc weight, Appendix S1: Table S2).

| CV | Best model | ED | R<sup>2</sup> | p |
| --- | --- | --- | --- | --- |
| Leaf production | Shannon index (H') |  | 0.14 | 0.22 |
| LAI | **Shannon index (H')** | + | **0.54** | **0.006** |
| Fruit production | Shannon index (H') |  | 0.17 | 0.18 |
| Stem increment | ECM trees [%] |  | 0.02 | 0.63 |
| Fine root biomass | **Shannon index (H') x ECM trees [%]** |  | **0.73** | **0.011** |
|  | Shannon index (H') | - |  | 0.007 |
|  | ECM trees [%] | - |  | 0.002 |
|  | Interaction | + |  | 0.010 |
| RAI | **Shannon index (H') x ECM trees [%]** |  | **0.71** | **0.015** |
|  | Shannon index (H') | - |  | 0.006 |
|  | ECM trees [%] | - |  | 0.003 |
|  | Interaction | + |  | 0.009 |

Plus and minus signs indicate the effect direction (ED). Statistically significant models are in bold.
LAI = leaf area index; RAI = root area index

### Influence of mycorrhizal type on tree biomass and surface areas

The proportion of ECM trees significantly influenced fruit production and LAI. The production of ECM fruits expedited total fruit production across the forest sites and increased distinctly with the proportion of ECM trees from 10 to 141 g m^-2^ a^-1^ (p < 0.001, R^2^ = 0.73; Fig. 1e). The production of fruits also increased with the increasing proportion of *F. sylvatica* (p = 0.01, R^2^ = 0.52). Concomitantly, the production of AM fruits decreased with the proportion of ECM trees from 14 to 5 g m^-2^ a^-1^ (p = 0.07, R^2^ = 0.28). The investigated ECM gradient also significantly influenced leaf surface area, increasing 75 % from a LAI of 4 m^2^ m^-2^ at higher pH sites with no ECM trees and low canopy height to a LAI of 7 m^2^ m^-2^ at lower pH sites dominated by tall ECM trees (p = 0.01, R^2^ = 0.55; Fig. 2c, Appendix S1: Figure S2c).

**Figure 1:**
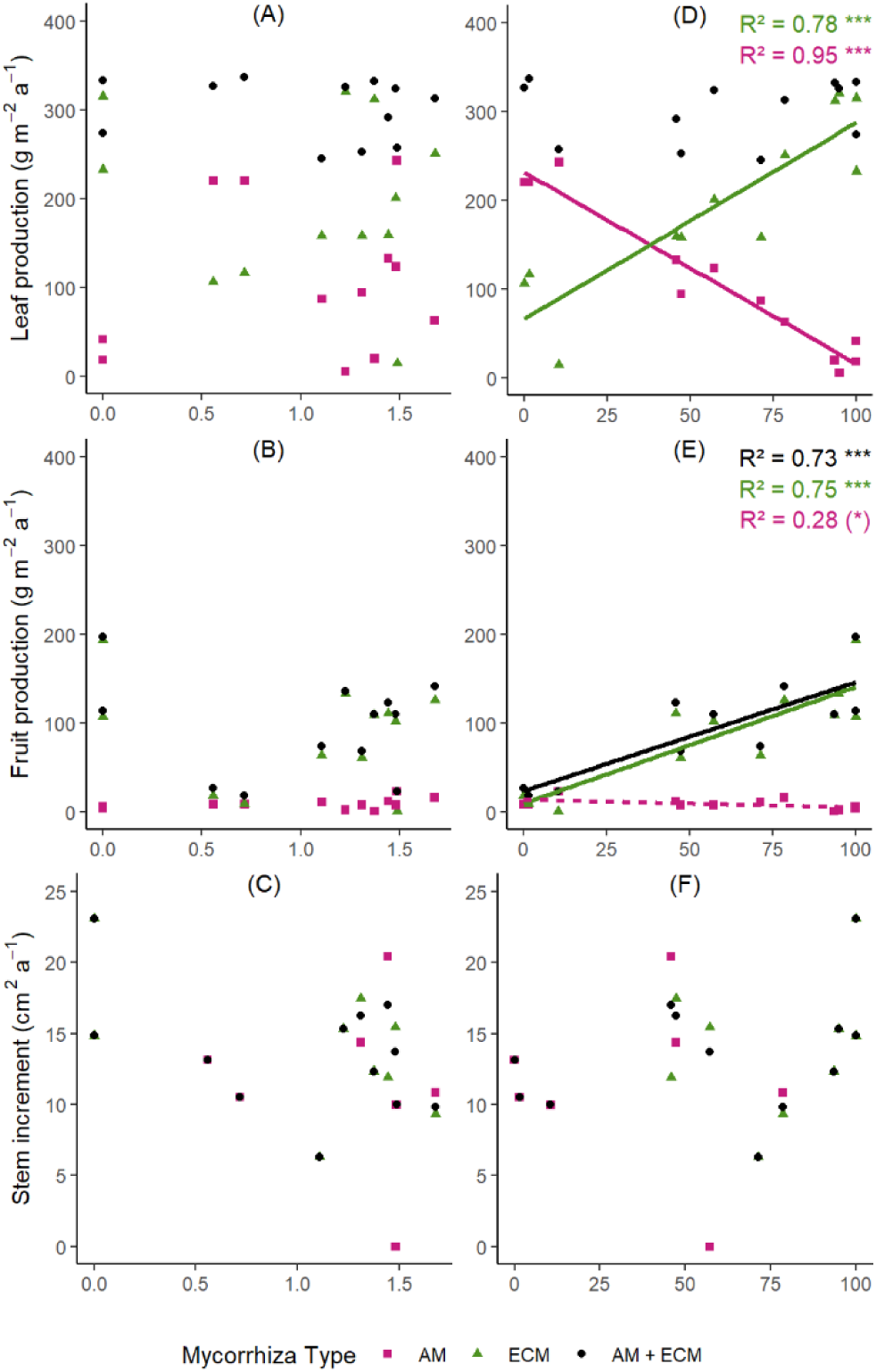
Influence of Shannon tree diversity (A to C) and the proportion of ectomycorrhizal (ECM) trees (D to F) on annual biomass production aboveground of arbuscular mycorrhizal (AM), ECM, and all trees of 12 mature deciduous forest sites. The influences on leaf production (A and D), fruit production (B and E), and wood production are shown as mean annual stem area increment of dominant trees (C and F) (Appendix S1: Table S3, S4).

**Figure 2:**
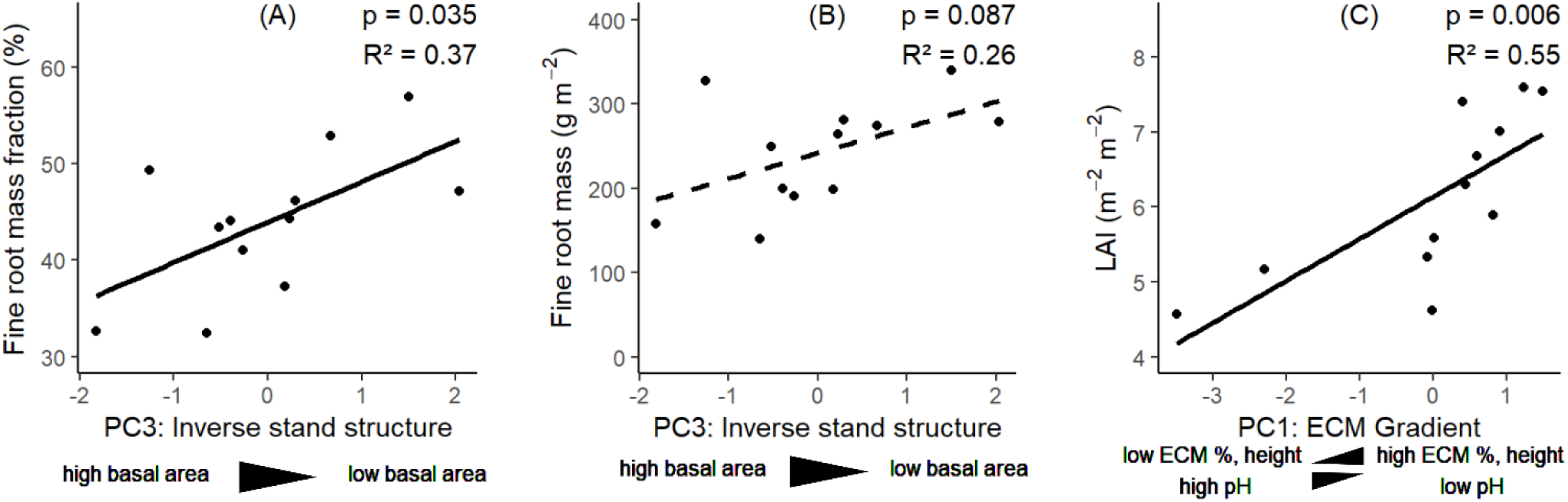
Effects of principal components (Table 2) on (A) fine root fraction (portion in fine root and leaf mass), (B) fine root mass, and (C) leaf area index (LAI) of 12 mature deciduous forest sites (Appendix S1: Table S5).

By contrast, the proportion of ECM trees did not influence stem increment (14 cm^2^ a^-1^ on average, Fig. 1f). It also had no significant influence on leaf mass (Appendix S1: Table S3), even though the production of ECM leaves significantly increased from 66 to 287 g m^-2^ a^-1^ (p < 0.001, R^2^ = 0.78; Fig. 1d) with an increase in the ECM trees from 0 to 100 % basal area, while the production of AM leaves decreased from 232 to 16 g m^-2^ a^-1^ (p < 0.001, R^2^ = 0.95; Fig. 1d). When ECM and AM trees had equal basal areas in the forest stand more ECM than AM leaves were produced (177 vs. 124 g m^-2^ a^-1^).

Belowground biomass was not influenced by the mycorrhizal association type. Fine root biomass averaged at 242 ± 127 g m^-2^ 20 cm^-1^ across all sites (Appendix S1: Figure S3b). Furthermore, the proportion of ECM trees did not affect the fine root surface area, which averaged at a RAI of 6 ± 3 m^2^ m^-2^ across all forest sites (Appendix S1: Figure S2d). Thus, biomass allocation to leaves and fine roots was not influenced by the proportion of ECM trees.

### Interaction of tree diversity and mycorrhizal type

The influences of tree diversity and mycorrhizal type on aboveground and belowground biomass and surface areas were mostly independent of each other. They only interacted with respect to the spatial variability of fine root biomass (p = 0.01) and RAI (p < 0.01; Table 3). The variability of both traits decreased with an increase in the proportion of ECM trees or an increase in Shannon diversity; that is, the higher the tree diversity or the more ECM trees were present, the more homogeneous fine root biomass and RAI were distributed in the investigated forest area. However, if the interaction was considered, the combined effect was less than the sum of the individual effects. That is, in cases where the proportion of ECM trees as well as the Shannon diversity were high, the positive effect on root distribution homogeneity was counteracted. In cases where both factors were low, the interaction effect was less pronounced.

### Influence of the stand structure on biomass allocation

Allocation of biomass was influenced by stand structure: More biomass was allocated into fine roots than into leaves (p = 0.04, R^2^ = 0.37), and fine root biomass was higher (p = 0.09, R^2^ = 0.26) when the basal area covered by trees decreased (Fig. 2a, b). Basal area did not influence aboveground biomass and surface areas (Appendix S1: Table S5).

## Discussion

Our gradient analysis in a temperate deciduous forest allows for the first time to systematically disentangle the effect of tree diversity, mycorrhizal association, and their cross-effects on biomass and surface areas of trees in a similar abiotic environment. It reveals that biomass allocation is neither influenced by tree diversity nor the mycorrhizal association, but is a function of basal area in this forest area, i.e., of forest structure. The responses of aboveground biomass and surface area to tree diversity and mycorrhizal association were independent of each other, while the spatial variability of fine roots was driven by their interaction. These results contradict a dominant control of the mycorrhizal association on tree diversity-productivity effects; both influences must be considered individually.

### Tree diversity does not enhance production

In contrast to our initial hypothesis, tree diversity did not increase aboveground production in the investigated mature deciduous forest. Considerable foundational biodiversity research in grasslands has demonstrated that a gain in species increases ecosystem productivity due to a close biodiversity-ecosystem functioning (BEF) relationship. The resulting increase in production has been proposed to be the consequence of either (i) a higher chance of including highly productive species (*selection effect*), (ii) facilitative interactions between species that promote growth, or (iii) increased complementarity in resource use in space and over time (*niche partitioning*). Yet so far, most BEF relationships have been inferred from grasslands or other non-forested ecosystems, while the consequences of tree species diversity for the functioning of mature forest ecosystems remain underexplored (Grossman et al. 2018) and partly inconclusive. Some evidence shows that mixed-species stands (often composed of two, sometimes more tree species) can be more productive in terms of biomass than monocultures with only one tree species (Zhang et al. 2012; Forrester and Bauhus 2016; Huang et al. 2018). A global meta-analysis showed that overyielding in old forests was dependent on environmental conditions and increased with higher precipitation (Jactel et al. 2018). Other studies contradicted these results (Meinen et al. 2009; Jacob et al. 2010; Seidel et al. 2013) and showed that stand structure can significantly affect BEF relationships in forest ecosystems and can be an even stronger determinant of productivity than tree diversity (Vilà et al. 2013; Paquette and Messier 2011). Positive BEF relationships in mature temperate and subtropical forest stands were identified when the effect of tree size was accounted for (Chamagne et al. 2017; Ren et al. 2021). This is confirmed with our study, in which higher basal area (of older or more dominant trees) was a determinant of biomass allocation to leaves – and not tree diversity. Many observational studies across tree diversity gradients employ dilution gradients, where one matrix species is always present, forming the baseline monoculture forest stand against which diversity effects are compared. This matrix species in temperate forests of Central Europe is often European beech, which is the dominant tree species in the natural forest vegetation and of economic importance (Bundesministerium für Ernährung und Landwirtschaft 2024). This is also the case in our gradient study, where sites with a Shannon diversity of 0 are monospecific beech forests, and sites of higher diversity are mostly mixed beech forests (except sites #3 and #11). European beech is a tall-growing, late-successional deciduous tree species that reaches dominance by casting deep shade with a high LAI (Leuschner and Meier 2018). A greater LAI promotes high photosynthetic capacity and thus productivity (Leuschner and Meier 2018; Ammer 2019; Reich 2012) and may be the reason for comparably high production in monospecific stands of our tree diversity gradient (Ratcliffe et al. 2015), which counteracts a selection effect. When Shannon diversity increases in mixed beech forests, beech has not reached dominance yet and is accompanied by other, earlier successional tree species of lower tree height and greater variability in LAI, as shown in our study. In deciduous Central European forests, the influence of European beech and tree diversity leads to contrasting influences on production that cancel each other out and are probably the reason for the inconclusive proof of overyielding in mature forest stands of Central Europe.

### Tree diversity influences the spatial variability of leaf and fine root areas

In the absence of overyielding, tree diversity increased the spatial variability of LAI, which reduced the density of leaf area packing. It has been demonstrated previously that old-growth beech forests with high species diversity have higher fine-scale variability in LAI than managed beech forests (Manes et al. 2010). Spatial variability of LAI can be explained by canopy gaps and standing dead trees in senescent forest growth stages, that is, the absence of foliage at local spots, which is crucial for tree regeneration and maintenance of species diversity. However, they can also result from the heterogeneity of canopy architectures and the functional diversity of crown allometry among different tree species, which leads to heterogeneous foliage. Increased spatial variability of LAI in diverse tree stands in our study opposes previous findings of denser-packed crowns in mixed-species forests, which is thought to support efficient exploitation of canopy space and maximize light interception (complementary effects; Jing et al. 2021). Yet crowns can also become more plastic as a result of intraspecific variation in crown morphology in response to competing neighbors (Seidel et al. 2011). Trees invest more C into height growth to compete for light while delaying their crown development (Lines et al. 2012), which leads to less canopy packing. Light competition between European beech and other tree species may be a reason for the greater spatial variability of LAI found in diverse forest sites of our study, where beech has not yet reached dominance. As a consequence, light transmission to the ground and nutrient availability may also be spatially more variable, which can support diversity of non-tree species in the canopy and herb layers of diverse forest stands (Dormann et al. 2020; Lewandowski et al. 2021) and enhance litter decomposition as a consequence (Jonsson et al. 2015).

Tree diversity and the proportion of ECM trees decrease the variability of fine root mass and RAI, resulting in more homogeneous root distribution in diverse as well as in ECM forest soils. The decrease in fine root variability in response to diversity is probably a result of more homogeneously occupied topsoil space at these sites, when niche partitioning from segregation into deeper, nutrient-poorer soil layers is restricted. This may also explain the absence of root biomass overyielding in diverse forests, as proposed by Wang et al. (2023), as the space for more root biomass is simply not available. ECM trees depend more on mycorrhizal hyphae when foraging into resource-rich soil spots than AM trees (Chen et al. 2018; Jevon and Lang 2022), which can lead to more homogenous root distribution in ECM forests, while AM root systems are more heterogeneous and used for exploration and subsequent exploitation. In addition, AM trees exhibit stronger inhibition by conspecifics than ECM trees (Bennett et al., 2017), which may contribute to greater non-uniformity in root distribution. Considering the interaction of both gradients, the positive effect of diversity and the proportion of ECM trees on root homogeneity is weaker when both are high, likely due to higher root heterogeneity in mixed ECM stands compared to pure beech stands. Thus, the direction of the effect of diversity on root homogeneity varies depending on the proportion of ECM trees and *vice versa*.

Taken together, tree diversity shifts spatial variability from belowground to aboveground biomass and surface area in the investigated forest, that is, leaf area becomes more heterogeneous, and root biomass and area become more homogeneous at diverse sites. This pattern cannot be explained by resource availability, which is homogeneous or directed aboveground and heterogeneous belowground, but is likely a result of competition processes and species-specific properties.

### ECM trees invest in light perception and reproduction

In partial agreement with our second hypothesis, LAI increased from AM to ECM forests, which supports light perception. LAI varies with the successional status of trees, generally increasing from early- to late-successional species (Leuschner and Meier 2018). In our study, focused on typical species of the Central European tree flora, AM trees represented early- to mid-successional stages, whereas ECM trees represented mid- to late-successional stages. This can explain the increase in LAI in ECM forests, probably particularly driven by hornbeam, European beech, and large-leaved lime, with a comparably high LAI of 7.1 to 8.3 m^2^ m^-2^ (Leuschner and Meier 2018). These species often colonize organic soils and acquire substantially more organic N than AM trees (Averill et al. 2014), which promotes the formation of a distinct shade crown and productivity but comes at the expense of greater frost-sensitivity in ECM forests composed of late-successional tree species.

The mycorrhizal association affected fruit production in our gradient study, where the increasing proportion of ECM trees increased fruit mass by a factor of 14, which can be a result of both individual fruits being heavier and/or more fruits being produced. Particularly, the ECM oak species have heavier fruit masses of on average 3.5 g per fruit, while the anemochorous AM ash and maple have light fruits of *c.* 0.1 g (Leuschner and Meier 2018). Fruit production in several (predator-dispersed) ECM trees, such as European beech and oak, is occurring at irregular periods of 5 to 8 years in the form of mast reproduction (Leuschner and Meier 2018), which protects species reproduction by over-satiating seed predators to let some seeds escape from consumption (Kelly and Sork 2002). These intervals of mast fruiting are increasing in frequency in recent years, triggered by higher total solar radiation during summer (Müller-Haubold et al. 2015). Mast fruiting in ECM trees comes at the expense of C, which is not available for other organs’ production during this time, resulting in lower production of wood and leaf mass in mast years and lower C concentration in the leaves (Mund et al. 2010; Müller-Haubold et al. 2015). This shift in C investment and litter composition is also likely to affect decomposition rates in forests dominated by mast-fruiting ECM trees. For beech cupules (woody phyllomes of high lignocellulose content), it was shown that the initial decomposition is similar to leaf litter decomposition, whereas the complete decay process is extended (Fukasawa et al. 2012). Consequently, higher fruit production through more frequent mast fruiting in ECM forests at the expense of leaf production decelerates the decomposition of litter and increases the retention time of C in the forest floor.

### Basal area increases leaf biomass allocation

We found that forests with higher basal area allocate more biomass into leaves than into fine roots. Basal area, which is a result of the cross-sectional area occupied by individual trees and tree density, is a common measure to describe aboveground forest growth. The positive relationship between basal area and leaf biomass allocation aligns with previous work that demonstrated that stand structure, and more specifically tree height, is a strong predictor of aboveground biomass allocation (Jevon and Lang 2022). According to the optimal partitioning theory, enhanced biomass allocation aboveground is a result of aboveground resource limitation, that is, a limitation of radiation energy for photosynthesis (McCarthy and Enquist 2007; Poorter et al. 2012). Forest density enhances light competition between individual trees, resulting in stem and leaf biomass allocation (Poorter et al. 2012; Chamagne et al. 2017; Jevon and Lang 2022). The relationship between basal area and leaf biomass allocation has consequences for the C residence time in soil. Root necromass causes more efficient C sequestration in soil than leaf litter, primarily by direct binding of depolymerized root-C to mineral surfaces without subsequent microbial transformation (Kaštovská et al. 2024). Such mineral-associated organic matter (MAOM) can persist in the soil for centuries to millennia (Witzgall et al. 2024; Witzgall et al. 2021; Lehmann et al. 2020). Higher leaf biomass allocation in forest stands with higher basal area will thus lead to less efficient soil C sequestration and faster C turnover.

In conclusion, our study shows that biomass allocation is a function of stand structure in mixed beech forests and not of tree diversity or mycorrhizal type. Moreover, the effects of tree diversity and mycorrhizal type on Central European forests are independent from each other and have to be considered individually. While tree diversity increases the spatial variability of aboveground biomass and surface area, greater proportion of ECM trees accelerates mast fruiting and promotes leaf expansion and the formation of a distinct shade crown. Previous research has shown that the maintenance of large leaf area in ECM beech forests is even decoupled from soil moisture conditions in summer (Meier & Leuschner 2008), which plays a key role in the survival strategy of late-successional ECM trees to cast deep shade, but puts them also at risk of physiological failures and C starvation in increasingly drier future conditions.

## Acknowledgments

The authors wish to thank Labiqa Zahid, Jónina Amelie Hummel, Charolina Riana Christianty, Tania Laraib Batool, and Birte Buske for helping with data collection. ICM wants to thank the Deutsche Forschungsgemeinschaft (DFG, German Research Foundation) for financial support awarded within the Heisenberg program (grant no. ME 4156/5-1). DY wants to thank the Next Generation Ecosystem Experiments-Tropics, funded by the U.S. Department of Energy (DOE), Office of Science, Office of Biological and Environmental Research, and the Oak Ridge National Laboratory managed by UT-Battelle, LLC, for the DOE under contract DE-AC05-1008 00OR22725. We thank the managers of the three Exploratories, Anna K. Franke, Miriam Teuscher, Robert Künast, and all former managers for their work in maintaining the plot and project infrastructure; Victoria Grießmeier for giving support through the central office, Andreas Ostrowski for managing the central database, and Markus Fischer, Eduard Linsenmair, Dominik Hessenmöller, Daniel Prati, Ingo Schöning, François Buscot, Ernst-Detlef Schulze, Wolfgang W. Weisser and the late Elisabeth Kalko for their role in setting up the Biodiversity Exploratories project. We thank the administration of the Hainich National Park for the excellent collaboration. The work has been partly funded by the DFG Priority Program 1374 “Biodiversity-Exploratories” (DFG-Refno.). Field work permits were issued by the responsible state environmental office of Thüringen.

## Author Contributions

ICM, DY, and AR designed and planned the research project. AR performed the research and analyzed the data. ICM, AR, and DY wrote the manuscript. All authors approved the final version of the manuscript.

## Conflict of Interest Statement

The authors declare no conflict of interest.

## Notes

### Competing Interest Statement

The authors have declared no competing interest.

